# The effects of data adequacy and calibration size on the accuracy of presence-only species distribution models

**DOI:** 10.1101/775700

**Authors:** Truly Santika, Michael F. Hutchinson, Kerrie A. Wilson

## Abstract

1. Presence-only data used to develop species distribution models are often biased towards areas that are frequently surveyed. Furthermore, the size of calibration area with respect to the area covered by the species occurrences has been shown to affect model accuracy. However, existing assessments of the effect of data inadequacy and calibration size on model accuracy have predominately been conducted using empirical studies. These studies can give ambiguous results, since the data used to train and test the model can both be biased.
2. These limitations were addressed by applying simulated data to assess how inadequate data coverage and the size of calibration area affect the accuracy of species distribution models generated by MaxEnt and BIOCLIM. The validity of four presence-only performance measures, Contrast Validation Index (CVI), Boyce index, AUC and AUCratio, was also assessed.
3. CVI, AUC and AUCratio ranked the accuracy of univariate models correctly according to the true importance of their defining environmental variable, a desirable property of an accuracy measure. Contrastingly, Boyce index failed to rank the accuracy of univariate models correctly and a high percentage of irrelevant variables produced models with a high Boyce index.
4. Inadequate data coverage and increased calibration area reduced model accuracy by reducing the correct identification of the dominant environmental determinant. BIOCLIM outperformed MaxEnt models in predicting the true distribution of simulated species with a symmetric dominant response. However, MaxEnt outperformed BIOCLIM in predicting the true distribution of simulated species with skew and linear dominant responses. Despite this, the standard performance measures consistently overestimated the performance of MaxEnt models and showed them as always having higher model accuracy than the BIOCLIM models.
5. It has been acknowledged that research should be directed towards testing and improving species distribution modelling tools, particularly how to handle the inevitable bias and scarcity of species occurrence data. Simulated data, as demonstrated here, provides a powerful approach to comprehensively test the performance of modelling tools and to disentangle the effects of data properties and modelling options on model accuracy. This may be impossible to achieve using real-world data.

## 1 INTRODUCTION

Presence-only data, i.e. point locations where a species has been recorded as being present, are frequently used in modelling the distribution of a species. They can be obtained from atlases, museum and herbarium records, incidental species observation databases and radio-tracking studies. These data have proven especially useful for widely distributed species that are prohibitive to systematically sample across their entire range (Solano & Feria 2007; Thorn *et al.* 2009). Recent progress in biodiversity informatics and development of online databases of biodiversity, i.e. Global Facility Information facility (GBIF), Atlas of Living Australia (ALA), LifeMapper, and National Institute of Invasive Species Science (NIISS), have contributed significantly to the accessibility of such data (Graham *et al.* 2004; Culham & Yesson 2011). However, data on the distribution of species is often biased (Rondinini *et al.* 2006; Newbold 2010; Anderson 2012). While highly accessible locations can be surveyed numerous times, remote locations are often poorly surveyed (Reddy & Dávalos 2003; Kadmon, Farber & Danin 2004).

The increasing use of presence-only data for modelling species distributions has been promoted by the development of modelling techniques that only require presence-only data. These methods include BIOCLIM (Nix 1986), DOMAIN (Carpenter, Gillison & Winter 1993), Ecological Niche Factor Analysis (ENFA; Hirzel *et al.* 2002), Support Vector Machine (SVM; Guo, Kelly & Graham 2005), and MaxEnt (Phillips, Anderson & Schapire 2006). Some of these methods have been implemented in the online biodiversity databases for non-experts to build species distribution models through user-friendly web interface (Jetz, McPherson & Guralnick 2012; Guisan *et al.* 2013). For example, BIOCLIM method is used to build species distribution models in LifeMapper and GBIF, while MaxEnt is used in ALA and NIISS.

As model prediction is considered invalid without a proper model evaluation process (Manel *et al.* 1999), several indices have been introduced to evaluate the accuracy of presence-only species distribution models. These include the Absolute Validation Index (AVI; Hirzel *et al.* 2006), Contrast Validation Index (CVI; Hirzel *et al*. 2006), Boyce index (Boyce *et al.* 2002; Hirzel *et al*. 2006), and indices based on the Receiver Operating Characteristic (ROC) plot (Hanley 1982). The area under the ROC curve (AUC), a commonly used index for assessing predictive performance of presence-absence models (Merckx *et al*. 2011), has been adapted for evaluating the accuracy of presence-only models by replacing absence data with background absences (Phillips, Anderson & Schapire 2006). Peterson, Papeş and Soberón (2008) later modified AUC to AUC_ratio_ by removing the need for background absences. Among these indices AUC and Boyce index are frequently used in presence-only distribution modelling, mainly because they circumvent the need for the identification of a specific threshold of performance (Franklin & Miller 2009).

Faced by a wide range of species distribution modelling methods, several studies have focused on comparing model predictive performance by means of finding the most superior models (Elith *et al*. 2006; Tsoar *et al.* 2007). These studies have found that the modelling methods do not perform equally in predicting species distributions and that each method can give different inferences about the important environmental determinants. However, comparing model accuracy on the basis on how well the model predicts the distribution of a species as reflected by the data, has been widely debated, as it can give ambiguous results regarding which models are best to predict species distributions (Zurell *et al.* 2010). This is mainly because the data that were used to train and validate the model can both be biased in representing the true distribution of a target species. The use of background or pseudo-absence data can also play a misleading role, because it may introduce an indeterminate number of false absences into models (Graham *et al.* 2004, Martínez-Meyer 2005).

Empirical studies can also give contradicting inferences about the impact of modelling options. For example, the choice of calibration size in the context of the characteristics of the species known occurrence data is an active research issue in modelling presence-only data (Van Der Wal *et al.* 2009, Webber *et al*. 2012, Merow, Smith & Silander 2013). Van Der Wal *et al.* (2009) found that the size of the calibration area impacts the accuracy of species distribution models derived using MaxEnt, and Thuiller *et al.* (2004) identified an optimal size of the calibration area based on an AUC accuracy index. On the other hand, Giovanelli *et al.* (2010) found MaxEnt to have limited sensitivity to calibration area.

With large compilations of presence-only data, a key challenge in assessing species distributions in conservation practice entails maximizing information return from limited data, i.e. developing and testing tools that are able to generate reliable results despite inevitable data bias and scarceness (Newbold 2010). However, in order to systematically perform such analyses, a comprehensive database is required that captures the nuances of species distributions (e.g. species rarity and detectability), the diverse ecological process that drives species distributions (e.g. species-environmental relationships, dispersal barriers) and the large variation of survey properties (i.e. survey resolution and extent) (Meynard & Kaplan 2012). Artificial species derived using simulated data provides a powerful way to capture this variation and have been used to assess the effects of various data characteristics, modelling options, and species distribution modelling tools (Zurell *et al.* 2010; Santika 2011; Meynard & Kaplan 2012). However, the application of this approach to assessing how the extent of survey data and the size of calibration area affect the performance of presence-only models has been limited to date.

Using simulated data, we addressed this limitation by focussing on three research questions: (1) how do presence-only indices perform in measuring the accuracy of presence-only models? (2) how does imperfect survey coverage data affect the accuracy of presence-only models? Finally, (3) how does calibration size affect the accuracy of presence-only models?, i.e. whether the study area tightly surrounds species presences versus more broadly capturing the species presences. We tested four presence-only accuracy indices, (CVI, Boyce index, AUC, and AUC_ratio_), against predictions derived from two modelling methods (BIOCLIM and MaxEnt).

## 2 MATERIALS AND METHODS

### 2.1 Simulated species data

The generation of simulated species data followed similar approach as described in Santika and Hutchinson (2009). The procedure is illustrated in Figure 1 with an example of one simulated species. First, ten environmental variables were simulated on a 100×100 square lattice with varying degrees of spatial autocorrelation. The variables were constructed so that they were not correlated with each other. Five of these variables were used to generate one simulated species (Fig. S1a), and the remaining variables were retained for modelling purposes only. We assumed that the simulated species responded in varying ways to each of the five environmental variables, i.e. symmetric, skew or linear. Various weights were assigned to each variable, reflecting the relative importance of each variable in determining the species distribution, with the sum of all five weights equal to one. A habitat suitability map of the species (Figure 1b) was then calculated as a sum of the products of these simulated variables and their assigned variable weights. We repeated this procedure 500 times using different combinations of environmental variables, variable weights and species-environmental responses, to produce a total of 500 habitat suitability maps.

**Figure 1.**
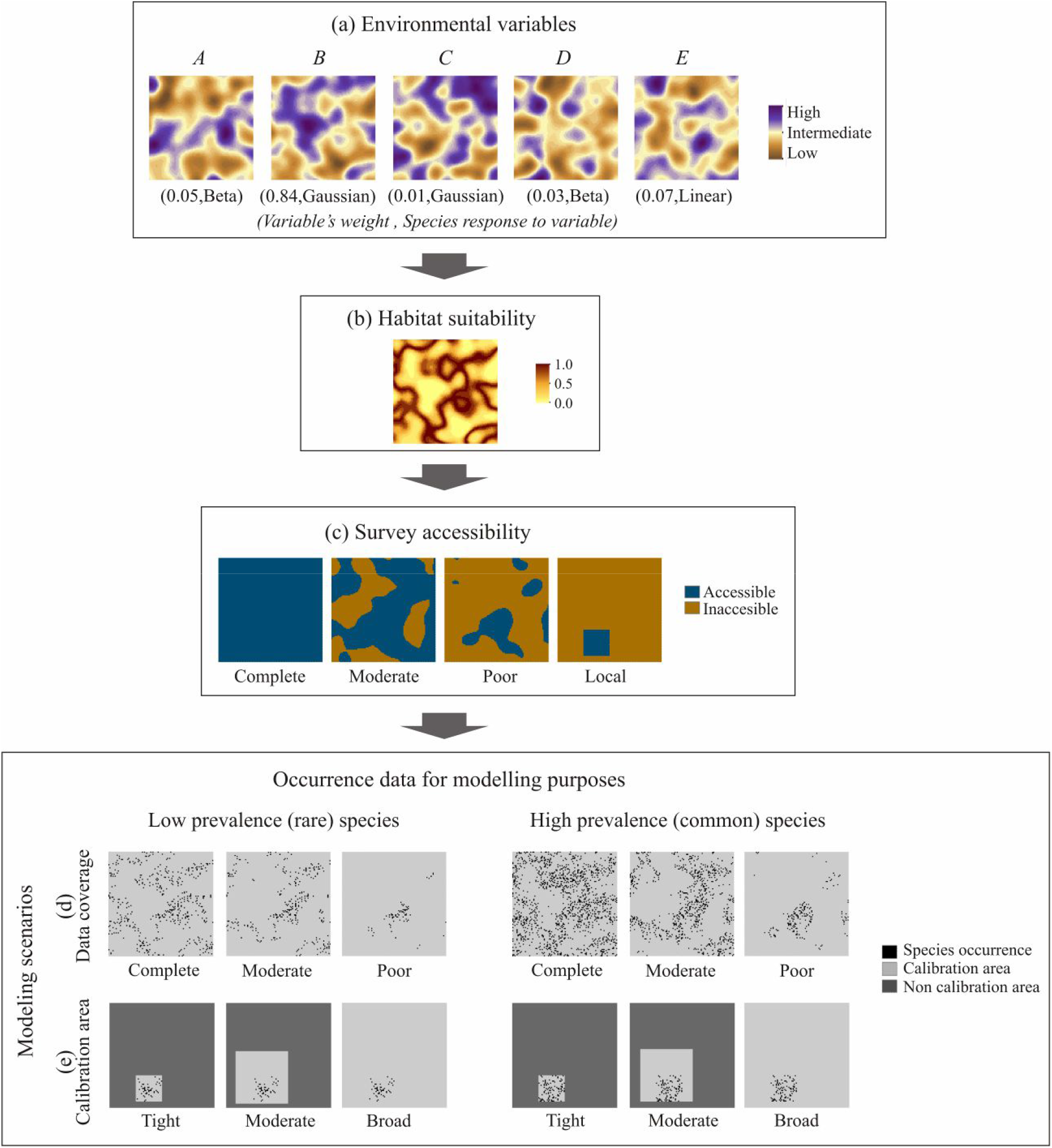
Schematic representation of steps required for generating one simulated species. (a) We first generated five spatially clustered environmental variables, and assigned a relative weight and type of species-environmental relationship response to each of these variables. (b) We derived a habitat suitability map based on these variables. (c) We generated four spatially clustered binary maps representing varying survey accessibilities. (d) We generated simulated species data with varying prevalence (low and high) and data coverage (complete, moderate and poor). (e) We generated local data and varied the calibration size with respect to the area covered by species occurrences (tight, moderate and broad).

Studies have shown that species prevalence (species being rare or common) crucially affects the accuracy of species distribution model (Manel, Williams & Ormerod 2001; Santika 2011). Taking into account this aspect, we derived two types of species for each habitat suitability map using a probabilistic approach (Meynard & Quinn 2007; Santika 2011): low prevalence (rare) species and high prevalence (common) species. Then for each type of species, we assumed three types of survey coverage data: complete, moderate, and poor (Figure 1d). Complete coverage data included the whole area of the 100×100 square lattice and represents complete distributional range of a species. The moderate and poor coverage data were obtained by multiplying the complete coverage data with a spatially clustered binary map representing some unknown factor supporting data collection effort (i.e. 1 denotes that a cell is accessible for data collection and 0 denotes that a cell is inaccessible) (Figure 1c). For the moderate coverage data, the spatially clustered binary map was obtained by applying a threshold of 0.5 to a spatially clustered map with continuous values between 0 and 1, so that values larger than or equal to 0.5 were assigned 1 and values smaller than 0.5 were assigned 0. The poor coverage data were defined by applying a threshold of 0.75.

Besides the three types of simulated data described above, we also generated a simulated dataset that includes only a subset area of the 25×25 square lattice (Figure 1c) of the complete survey data to reflect varying calibration sizes on model accuracy, i.e. tightly surrounding species presences (encompassing only the 25×25 square lattice), moderately surrounding species presences (encompassing a 50×50 square lattice around the local dataset), and broadly surrounding species presence data (encompassing the entire 100×100 square lattice around the local dataset) (Figure 1e). A total of 4,000 simulated species datasets were generated (500×2×4).

### 2.2 Modelling scenarios

Ten univariate models (using five relevant variables and five irrelevant variables) were fitted to each of the 4,000 simulated species with varying extents of survey data (i.e. complete, moderate, and poor survey) and prevalence using BIOCLIM and MaxEnt. BIOCLIM predictions were generated using DIVA-GIS software (Hijmans *et al*. 2005). MaxEnt predictions were generated using MaxEnt software (version 3.4.0) (Phillips, Anderson & Schapire 2006) using the default settings. Model predictions were generated for the entire 100×100 square lattice. First, we used the models generated using the complete dataset to assess the validity of presence-only evaluation measures in correctly identifying important environmental determinants. Second, we used the models generated using the three different datasets to assess how survey coverage, i.e. complete, moderate, and poor coverage, affects model accuracy. Finally, we fitted the ten univariate models to each of the 1,000 local datasets to vary the size of calibration area. This represents a scenario whereby species survey data is only available locally and covers only a small portion of the entire species distributional range.

### 2.3 Modelling method evaluation

For each species distribution model prediction, we conducted model evaluations using four accuracy indices: CVI, Boyce index, AUC, and AUC_ratio_, and against two types of data. First, each prediction was evaluated against the complete survey data (100×100 square lattice), to measure the accuracy of a model in predicting the true distribution of a species (in practice this distribution is likely to be unknown). Second, each prediction was evaluated against the same set of data used for model training, to measure the accuracy of the model in predicting the apparent distribution of a species as reflected by the training data. This is the modelling measure normally available in practice. To differentiate between model predictive performance based on these two approaches, we added a subscript *(true)* and *(model)* after each evaluating measure to denote the evaluation result that is based on the first approach (true accuracy) and second approach (model accuracy), respectively. For example, AUC_ratio*(true)*_ and AUC_ratio*(model)*_ denotes the true accuracy and model accuracy, respectively.

CVI is calculated as the proportion of presence points falling in cells having a threshold habitat suitability index minus the proportion of cells within this range of threshold of the model (Hirzel *et al.* 2006). Because the value of CVI varies depending on the choice of threshold, we calculated the model’s accuracy based on CVI using varying thresholds, i.e. CVI_0.25_ (i.e. CVI with threshold 0.25), CVI_0.5_, and CVI_0.75_.

The calculation of Boyce index requires the habitat suitability range to be partitioned first into *b* classes (or bins) (Boyce *et al.* 2002). For each class *i*, we calculated the predicted frequency of evaluation points, *P*_*i*_, and the expected frequency of evaluation points, *E*_*i*_. *P*_*i*_ was calculated as the number of evaluation points predicted by the model to fall in the habitat suitability class *i*, over the total number of evaluation points. *E*_*i*_ was calculated as the number of grid cells belonging to habitat suitability class *i*, over the total number of cells in the whole study area. The Boyce index was then calculated as a Spearman correlation between *P*_*i*_/*E*_*i*_ and class *i*. Here, we calculated the value of Boyce index for various bin sizes, i.e. Boyce_bin(10),_ Boyce_bin(20)_ and Boyce_bin(50)_.

The main shortcoming of the bin-based Boyce index is its sensitivity to the number of suitability classes *b* and to their boundaries (Boyce *et al.* 2002). Hirzel *et al.* (2006) have proposed a way to fix this problem by deriving a new evaluator based on a moving window of width *W*, instead of fixed classes. Computation begins with the first class covering the suitability range [0,*W*] whose *P/E* ratio is plotted against the average suitability value of the class, *W*/2. Then, the moving window is shifted from a small amount upwards and the *P*/*E* is plotted again. This operation is repeated until the moving window reaches the last possible range [1-*W*, 1]. This provides a smooth *P*/*E* curve, in which a continuous Boyce index, Boyce_cont(*W*_) is computed. We tested varying window sizes, i.e. Boyce_cont(0.1)_, Boyce_cont(0.2)_ and Boyce_cont(0.5)_.

AUC was calculated as the area under the Receiver Operating Characteristic (ROC) curve. A ROC curve is obtained by plotting all sensitivity values (the proportion of correctly predicted presence) on the *y*-axis against 1-specificity (1 minus the proportion of correctly predicted absence) values on *x*-axis for all available thresholds. Phillips, Anderson and Schapire (2006) adapted the AUC approach for measuring discriminatory power of MaxEnt models by replacing the absence with background data. In the absence of real absence data, we generated background samples as many as the number of presence cells used for the calculation of AUC. Peterson, Papeş and Soberón (2008) later modified the conventional AUC to be used for evaluating presence-only models, by removing entirely of the use of background data. Instead of plotting the sensitivity against the value of 1-specificity, the modified ROC curve plots the proportion of presences falling above a range of thresholds against the proportion of cells falling above the range of thresholds. The area under the modified ROC curve was then called AUC_ratio_.

## 3 RESULTS

### 3.1 The performance of evaluation measures

There is a strong correlation between the accuracies of MaxEnt univariate models (generated using the complete survey data) based on AUC_*(true)*_, AUC_ratio*(true)*_ and CVI_0.5*(true)*_ and the weight of the variables (Pearson correlation *r*P ≥ 0.9, Figure 2). Similar results were obtained for CVI_0.25*(true)*_ and CVI_0.75*(true)*_, and for the BIOCLIM univariate models. This indicates that the AUC, AUC_ratio_ and CVI were able to rank the accuracies of univariate models based on the importance of their defining environmental variable. Species with low prevalence were rated higher than species with high prevalence based on these measures. However, this was not the case for Boyce index. The accuracies of MaxEnt univariate models based on Boyce_cont(0.2)*(true)*_ and Boyce_bin(10)*(true)*_ was poorly correlated with the variable’s weight (*r*P ≤ 0.25). Similar results were obtained for Boyce_cont(0.1)*(true)*_, Boyce_cont(0.5)*(true)*_), Boyce_bin(20)*(true)*_ and Boyce_bin(50)*(true)*_, and for the BIOCLIM univariate models. This indicates that Boyce index failed to rank the accuracies of the univariate model correctly. Furthermore, while models generated using variables that were irrelevant for the distribution of the simulated species had mostly very low AUC_*(true)*_, AUC_ratio*(true)*_ and CVI_*(true)*_, 73% of these models had Boyce_cont(0.2)*(true)*_ higher than 0.8. High percentages (67% and above) were also observed in other Boyce indices for these models.

**Figure 2.**
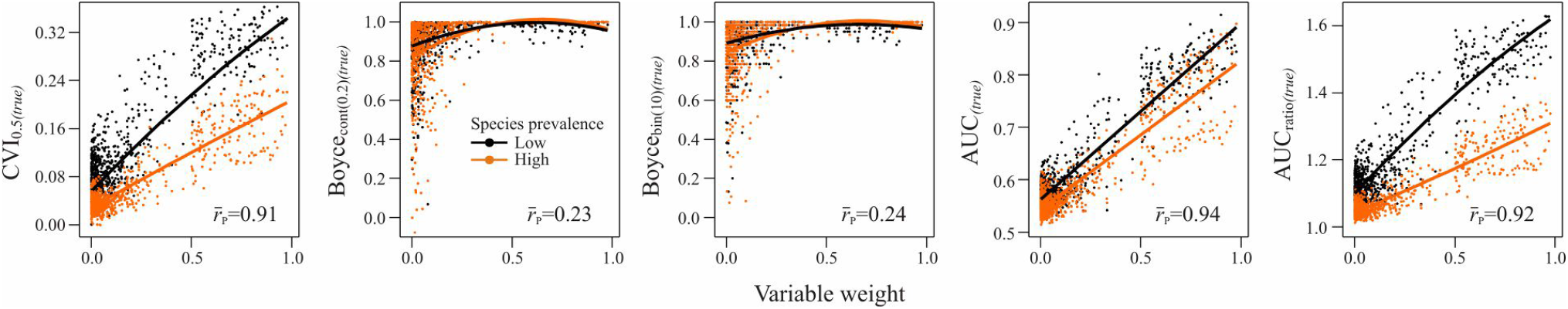
The relationships between the accuracy of MaxEnt univariate models, based on CVI_0.5*(true)*_, Boyce_cont(0.2)*(true)*_, Boyce_bin(10)*(true)*_, AUC_*(true)*_, and AUC_ratio*(true)*_, and the variables’ weights representing the relative importance of the variables in determining the distributions of the simulated species, by species prevalence. 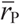 denotes the mean Pearson correlation of the relationships across simulated species with low and high prevalence. Similar results were obtained for the BIOCLIM models.

### 3.2 The effect of data adequacy and calibration area on model performance

Inadequate data and wide calibration areas hampered the identification of the most important environmental variable for a species. The proportion of environmental variables correctly identified as the most important by both BIOCLIM and MaxEnt methods, based on AUC_ratio*(model)*_ decreased if the data was poorly sampled across the species range or as the size of the calibration area increased (Figure 3a-b). However, the ability of the modelling methods to identify correctly the most important variables for a species under varying data adequacies and calibration areas depended on the type of species-environmental response. BIOCLIM typically outperformed MaxEnt for identifying correctly the most important variables with a symmetric or skew response. In contrast, MaxEnt outperformed BIOCLIM for identifying correctly the most important variables with a linear response. The performance of MaxEnt method improved across all species-environmental relationships with complete data. Similar results were obtained for AUC_*(model)*_ and CVI_*(model)*_.

**Figure 3.**
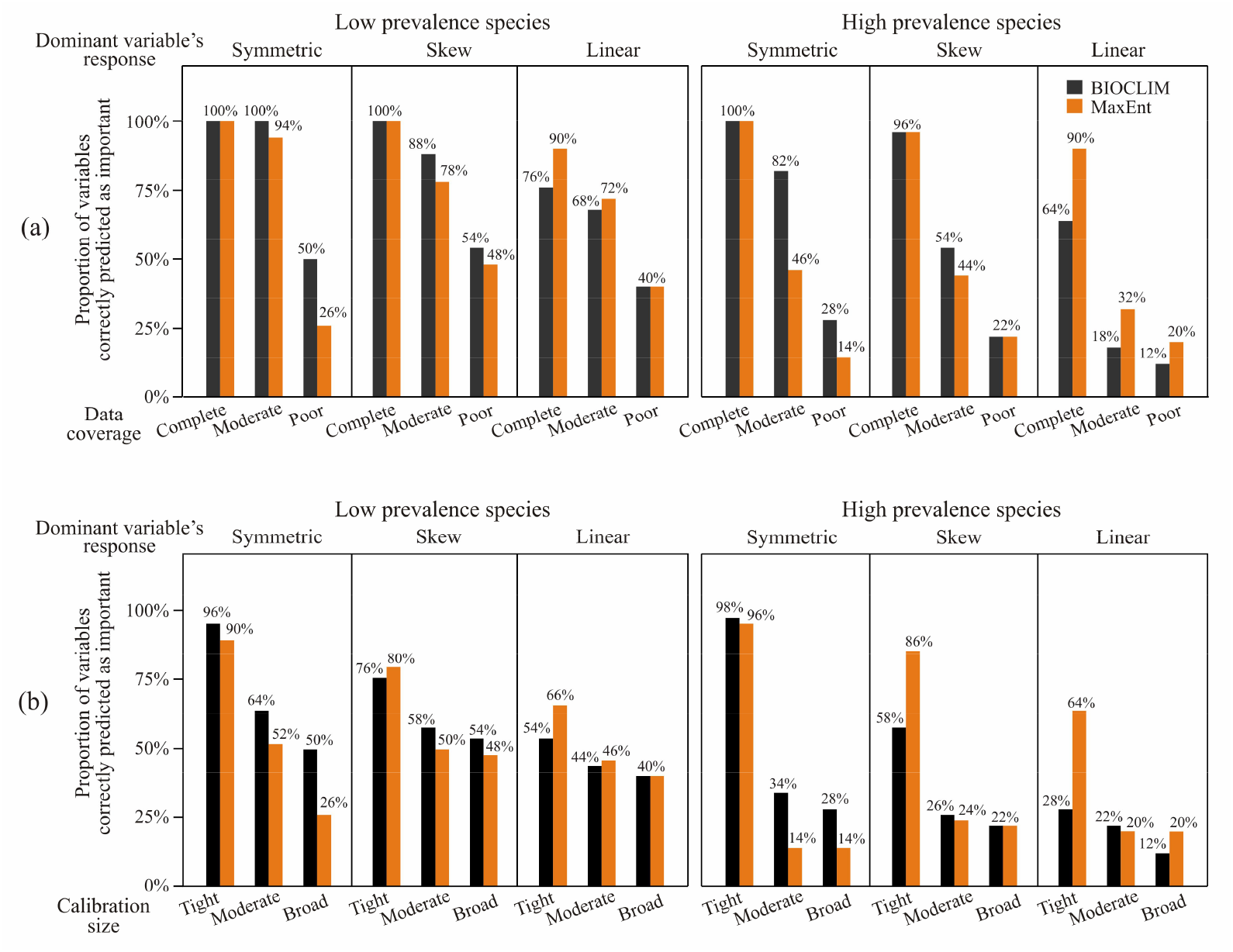
The proportions of variables correctly predicted as the most important by BIOCLIM and MaxEnt, by species dominant response and prevalence, given (a) varying data coverage (complete, moderate, or poor) and (b) varying calibration size with respect to the area covered by species occurrences (tight, moderate, or broad).

Consequently, when BIOCLIM and MaxEnt univariate models, based on variable identified as the most important, were evaluated against the true simulated species data, the accuracies of the models based on AUC_ratio*(true)*_ reduced if data was poorly sampled (Figure 4a) or as the calibration area increased beyond the area capturing species presences (Figure 5a). However, when the models were evaluated against imperfect training data via cross validation, the discriminative power of the model based on AUC_ratio*(model)*_ increased if the data was poorly sampled (Figure 4b) or as the calibration area increased beyond the area capturing species presences (Figure 5b). Interestingly, despite the fact that BIOCLIM models were slightly more accurate in predicting the true distribution of species with a symmetric dominant response than MaxEnt models (due to the number of variables correctly predicted as important were higher for BIOCLIM than for MaxEnt), the discriminative power of MaxEnt models were consistently higher than BIOCLIM models. The ratio between the discriminative power and the accuracies of the univariate model provides a measure of how far the model performance strays from the truth as the quality of data reduced. Here we found that AUC_ratio*(model)*_/AUC_ratio*(true)*_ was higher for MaxEnt than for BIOCLIM, suggesting MaxEnt overestimated the model performances compared with BIOCLIM under limited data coverage (Figure 4c) and broad calibration areas (Figure 5c). Similar results were obtained for AUC and CVI.

**Figure 4.**
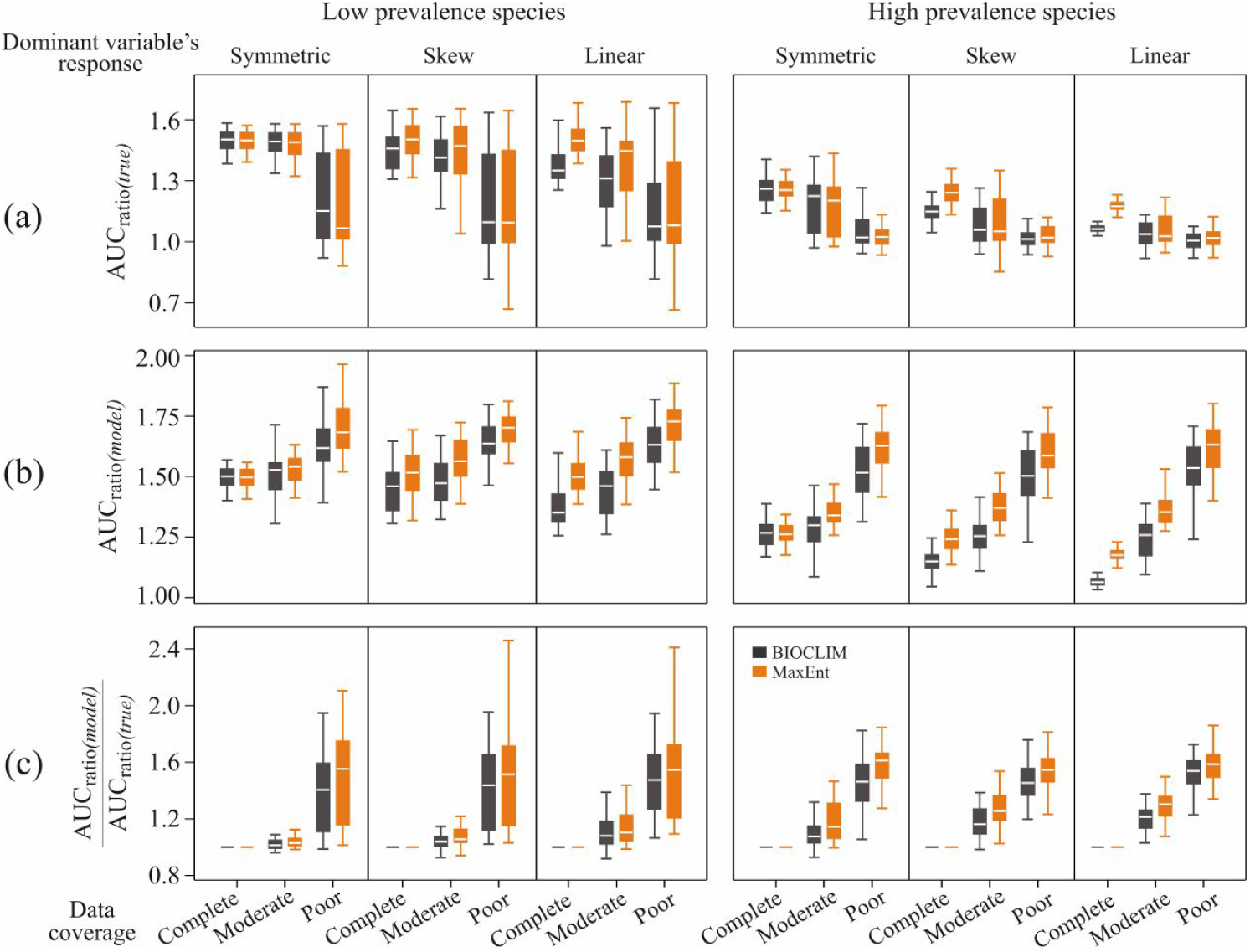
The effect of varying data coverage (complete, moderate or poor) on (a) models’ true accuracy based on AUC_ratio*(true)*_, (b) models’ accuracy based AUC_ratio*(model)*_, and (c) the ratio between models’ accuracy and the true accuracy (AUC_ratio*(model)*_/ AUC_ratio*(true)*_) as a measure of model inflation factor, as generated by BIOCLIM and MaxEnt methods

**Figure 5.**
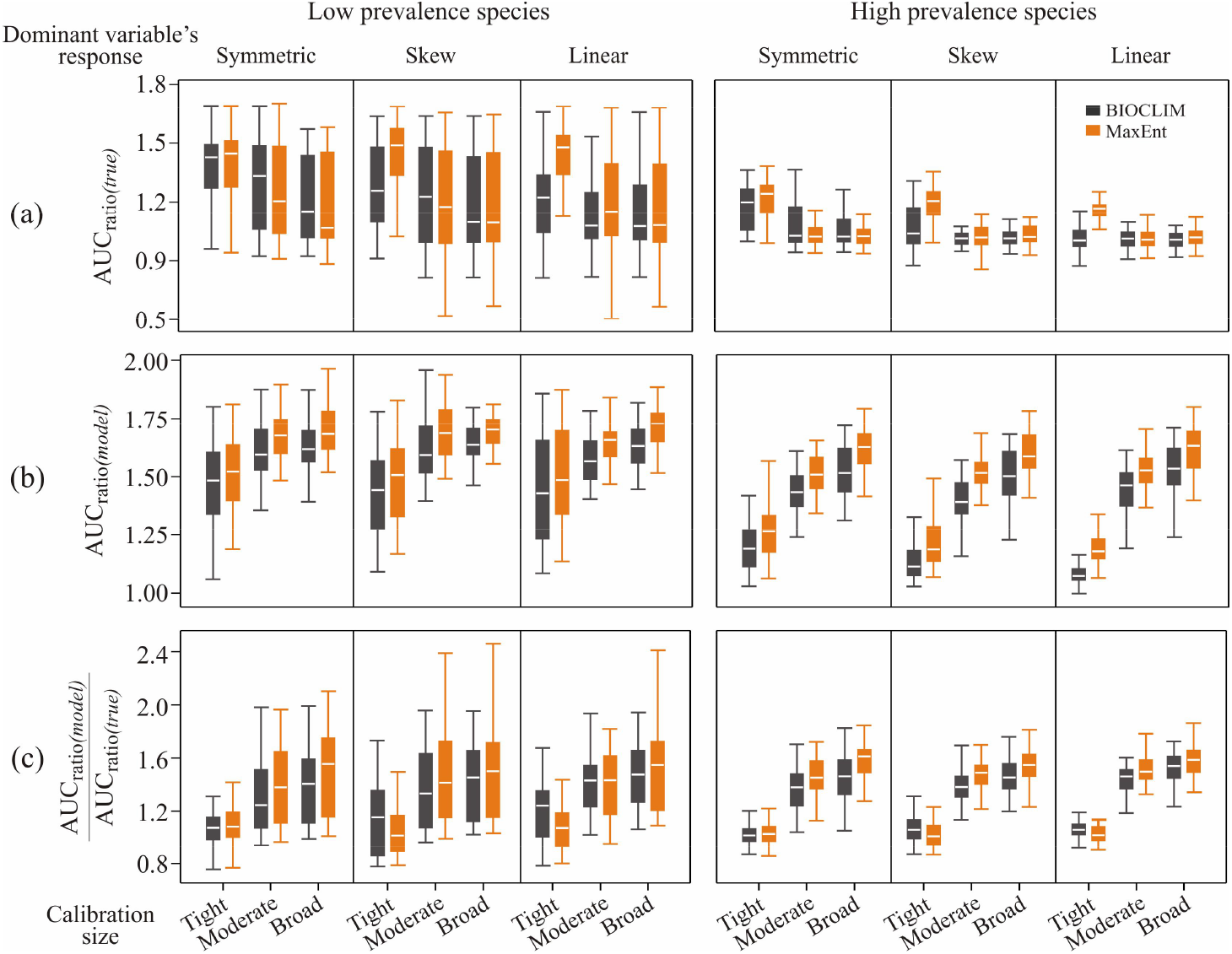
The effect of varying calibration size with respect to the area covered by species occurrences (tight, moderate or broad) on (a) models’ true accuracy based on AUC_ratio*(true)*_, (b) models’ accuracy based AUC_ratio*(model)*_, and (c) the ratio between models’ accuracy and the true accuracy (AUC_ratio*(model)*_/ AUC_ratio*(true)*_) as a measure of model inflation factor, as generated by BIOCLIM and MaxEnt.

## 4 DISCUSSION

Many modelling methods and evaluation measures have been developed to accommodate presence-only species distribution data (Nix *et al.* 1986; Hirzel *et al.* 2006; Phillips, Anderson & Schapire 2006). However, the assessment of these tools has mostly been based on empirical data (Elith *et al.* 2006; Liu *et al.* 2005). Since the data used to train and test model accuracy is often biased, inference regarding the performance of these tools can be unreliable (Zurell *et al.* 2010; Yackulic *et al*. 2013). As demonstrated here, simulated data provide a way to assess comprehensively the accuracy of the modelling tools as well as aspects of the data and model options. This may not be possible to achieve using real-world data.

Here we found that the Boyce index method of model evaluation performed poorly when ranking the accuracy of univariate models based on the variables’ relative importance. Furthermore, high proportions of irrelevant variables yielded MaxEnt univariate models with high Boyce index. This implies that there is a high chance that variables which are not relevant in determining the distribution of a species and would provide an unreliable prediction map, would appear superior based on Boyce index. Our finding concurs with previous studies that reported relatively high model accuracies based on the Boyce index while other indices showed relatively low model accuracies (Sattler *et al*. 2007; Braunisch & Suchant 2010; Bellard *et al*. 2013). This is of concern, considering that Boyce index is commonly used to assess the performance of species distribution models. A literature search in Google Scholar (10 November 2017) shows that Boyce index has been used in 92 empirical studies published between 2007 and 2017, and was the second most popular measure after AUC, while AUC_ratio_ (15 papers) and CVI (24 papers) were used less frequently. Furthermore, many of these studies have used the Boyce index alone to assess model accuracy (Galparsoro *et al.* 2009; De Angelo Paviolo & Di Bitetti 2011; Petitpierre *et al*. 2012; Rutishauser *et al.* 2012; Pocewicz *et al.* 2013). Our study shows that the use of Boyce index in modelling species distribution should be considered cautiously and that other accuracy measures, such as CVI, AUC and AUC_ratio_ should be given preference.

The dependency of accuracy measures on species prevalence has been examined previously, but mainly within the context of presence-absence species distribution modelling (Santika 2011; Meynard & Kaplan 2012). Here we have demonstrated that presence-only accuracy measures, such as CVI and AUC_ratio_, are also not immune to this species characteristic. Similar to the conventional AUC measure, both CVI and AUC_ratio_ tended to rank models predicting species with low prevalence (rare) more highly than models based on species with high prevalence (common) (Santika 2011). On the other hand, Boyce index was not affected by prevalence, perhaps due to its apparent lack of effective discrimination. The sensitivity of CVI on prevalence and the insensitivity of Boyce index on prevalence has also been reported by Hirzel *et al.* (2006) based on non-forest mountain plants distributional data in Switzerland.

Another advantage of assessments using simulated data is that it enables extensive exploration of why some modelling methods perform better than others (Elith & Graham 2009). This study shows that MaxEnt outperforms BIOCLIM mainly due to the flexibility of MaxEnt algorithm in fitting various species-environmental response functions, including the symmetric, skew and linear responses (Phillips, Anderson & Schapire 2006). However, BIOCLIM and MaxEnt had the same performance when predicting species with a dominant symmetric response. This is mainly because the BIOCLIM algorithm assumes that the probability of species occurrence is high as the environment is closer to the core (or median) of the species environmental range and declines gradually to zero at the limits of the range (Nix *et al.* 1986). BIOCLIM models therefore tend to be more effective when the species have been sampled from the full environmental range rather than being artificially localised, such as within a particular elevation range. Even in such cases, symmetric responses can be dominant (Oksanen & Minchin 2002). However, since in reality many other types of species-environmental responses exist, particularly when the survey area is limited, e.g. linear, skew, truncated (Austin 2002), the inability of BIOCLIM to fit other response functions is considered as a major drawback of this method. Furthermore, the BIOCLIM algorithm has no variable selection procedure, hence it assumes that each environmental predictor contributes equally in shaping the distribution of a species (Hijmans & Graham 2006). These factors explain why BIOCLIM generally perform poorly in distribution modelling studies using real-world species data and multiple environmental predictors (Elith *et al.* 2006).

There has been a common consensus that species distribution modelling research should be directed towards developing robust methods that are able to deal with the inevitable bias and scarcity of species occurrence data (Graham *et al.* 2004; Newbold 2010). Many studies have followed this guideline, attempting to find the best method that is able to predict species distribution given various real-world data conditions (Elith *et al.* 2006; Tsoar *et al.* 2007). MaxEnt methods have often been shown to produce models with highest model accuracy among other methods in species distributional modelling assessments (Elith *et al.* 2006). However, our simulation study has shown that the performance of MaxEnt is not as superior as standard performance measures would suggest, particularly in relation to BIOCLIM. Focussing only on simulated species with dominant symmetric responses (in which both MaxEnt and BIOCLIM methods performed equally well), we found that BIOCLIM models performed better than MaxEnt models in terms of providing correct inference about the dominant environmental variables and in predicting the true distribution of the simulated species, even in the context of inadequate data coverage and varying calibration sizes. Despite the fact that BIOCLIM models had higher true accuracies than MaxEnt models in predicting the distribution of the simulated species, MaxEnt models consistently outperformed BIOCLIM in predicting the apparent distribution of the species as reflected by the training data. This suggests that the true accuracy of MaxEnt models has been overrated, especially when there has been a lack of data coverage within a large modelling extent. Bean, Stafford & Brashares (2012) has found similar results on how small sample size affected the performance of MaxEnt models. Using limited data of the giant kangaroo rat (*Dipodomys ingens*) in California, USA, the authors found that the accuracy of MaxEnt models appeared to be much higher than it would have been if it had been evaluated against more complete data of the species’ distribution.

Our study also found that, with lack of complete knowledge regarding a species distributional range, correct identification of important environmental variables and accurate prediction of the species’ distribution can be maximized by confining the calibration area to tightly surround the species’ occurrence data. This supports the recommendation made by Elith *et al.* (2011). This applied to both BIOCLIM and MaxEnt models. Merow, Smith and Silander (2013) provides recommendations about how to choose calibration area for MaxEnt models. They recommend that the calibration area should be chosen depending on the study aim: if one were interested in determining the best location for a reserve for a species, then the background should be drawn only from the species known range, whereas if one were interested in which regions the species might invade worldwide in the absence of dispersal limitation, then the background should be chosen across the globe. However, applying a substantially large calibration area beyond the known species occurrences, as noted in the authors’ second recommendation, may increase the number of false background-absences and therefore yield a high model accuracy (Van Der Wal *et al.* 2009; Dupin *et al.* 2011). This would however also mask the true model accuracy and may result in erroneous under-prediction of the species’ potential distribution (Peterson, Papeş & Eaton 2007). This can be critical for the case of invasive species, as underestimating the species potential distribution can lead to an incorrect decision to allow import of species (Pheloung, Williams & Halloy 1999), which might subsequently lead to high mitigation costs (Yokomizo *et al.* 2009). Underestimation of species potential distribution can also lead to invaders being incorrectly labelled as harmless, leading to failure to establish early surveillance measures (Guisan *et al.* 2013).

This study demonstrated the power of using simulated data to assess the performance of species distribution modelling tools and to disentangle the confounded effects of species characteristics, species ecological process, sample properties, and modelling options on model performances. Our approach provides a significant advance over previous studies based on empirical data. This approach enables us to make a fair comparison among different modelling tools (both modelling methods and evaluation measures), and to better understand why some tools perform better than others. Understanding the strengths and limitations of a modelling tool is imperative, since it can inform users on how to apply the tool cautiously in the context of particular data structures and for particular species, as well as provide guidance on future research priorities for tool development.

## ACKNOWLEDGEMENTS

We acknowledge the Australian Government’s National Environmental Research Program and the Australian Research Council Centre of Excellence for Environmental Decisions for funding and support. KAW was supported by an Australian Research Council Future Fellowship.

